# How flat is your sample? An opportunistic survey of 3D tilt in public fluorescence microscopy data

**DOI:** 10.64898/2026.05.21.726891

**Authors:** Jacques Brocard

## Abstract

Sample planarity is rarely monitored in fluorescence microscopy quality control, yet focal plane deviations across the field of view are a potential source of measurement error. Here I describe FlatStat, a tool that estimates sample tilt automatically from any 3D fluorescence stack, without prior knowledge of sample content, by fitting a plane to the Z-map of maximum intensity. Applied to an Argolight calibration slide and biological samples on a laser-scanning confocal system, FlatStat yielded reproducible slope and direction measurements attributable to the instrument rather than the sample. To establish community reference values, FlatStat was extended to Python and applied opportunistically to 1204 image stacks from 22 projects in the Image Data Resource, yielding 4670 tilt measurements. Slopes spanned several orders of magnitude across projects; inter-channel coherence confirmed that measured tilt reflects physical stage and mounting geometry rather than channel-specific biological topography. Unfortunately, instrument and sample preparation metadata were largely absent from the corpus, limiting causal inference. Finally, controlled tilt experiments on fluorescent beads showed that chromatic shift increased modestly with tilt (∼57 nm over the full range tested), while lateral and axial resolutions were essentially unaffected.

## Introduction

Running a microscopy facility means occasionally fielding complaints. One day, a user reported that her samples looked “not flat” on one of our core facility laser-scanning confocal systems. A quick look confirmed the observation, something was indeed slightly off with the focal plane across the field of view. The most straightforward explanation was offered: perhaps the biological sample wasn’t flat or the coverslip mounting was suboptimal.

However, lingering doubts prompted more systematic investigations. How flat are our stages versus our samples? And how would any residual tilt affect precise 3D measurements? Searching the literature for answers proved surprisingly unrewarding. Standard quality control frameworks for fluorescence microscopy - including the widely adopted MetroloJ_QC suite (Faklaris *et al*., 2022) - make no mention of sample planarity as a monitored parameter. The only study I could identify that directly addresses the optical consequences of a tilted sample in fluorescence microscopy concerns a highly specific context: coverslip tilt in *in vivo* STED and two-photon imaging through a cranial window (Le Bourdellès *et al*., 2024), itself building on earlier limited work on water-immersion objective aberrations. Whether analogous effects occur in routine confocal imaging of conventionally mounted samples, and how prevalent such tilt is across the community, remained entirely open questions.

Here I describe FlatStat, a tool for automated planarity measurement from any 3D fluorescence stack, initially implemented as an ImageJ macro. Faced with the absence of community reference values against which to interpret our measurements, I extended FlatStat to Python and applied it opportunistically to survey a large corpus of publicly available fluorescence microscopy data from the Image Data Resource (IDR). Finally, returning to the confocal system that prompted this investigation, I quantified the optical consequences of a significant tilt on lateral and axial resolutions and chromatic shift using fluorescent beads.

## Methods

### FlatStat algorithm

FlatStat estimates sample planarity from any 3D fluorescence stack without prior knowledge of the sample content. The stack is downscaled by bilinear interpolation to a working resolution of approximately 100 pixels along its short axis. For each pixel, the Z position of maximum intensity is recorded across the stack, yielding a Z-map — a spatial map of the focal plane. A best-fit plane is then estimated by least-squares regression over the brightest pixels of this Z-map (top 10% of the max-intensity projection). The slope magnitude is expressed in µm/100µm, and the tilt direction as a compass bearing (0–360°). FlatStat is available as an ImageJ macro tool “FLATSTAT_v8.ijm” at https://github.com/jbrocardplatim/FLATSTAT and *doi:10.5281/zenodo.20325392*

To enable large-scale analysis, FlatStat was reimplemented in Python (“idr_flatstats_v6.py”, same repository). The Python implementation introduced two modifications relative to the ImageJ macro: the analysis was restricted to a central subset of N planes (N = min(50, max(5, ⌈2.5/p_z⌉)), with p_z the axial pixel size in µm), ensuring coverage of at least 2.5 µm regardless of acquisition settings; and the regression was performed independently at each of five percentile thresholds (88th to 92nd) of the max-intensity projection, providing a stable selection of bright, well-focused regions. The reported slope and direction are the mean and standard deviation across these five estimates.

### IDR corpus

The Image Data Resource (IDR, idr.openmicroscopy.org; Williams et al., 2017) was surveyed exhaustively using a custom Python script interfacing with the OMERO API (Open Microscopy Environment, 2023). Of 144 projects indexed at the time of the survey, eligibility was assessed on the basis of four criteria: imaging method containing “confocal” or “fluorescence” (excluding light-sheet, SPIM and related modalities); stack depth SizeZ > 5; minimum field dimension min(SizeX, SizeY) > 100 pixels; and availability of XY pixel size metadata. Projects exceeding 10,000 images were excluded to avoid corpus domination by high-content screening datasets. Up to 5 image stacks per dataset were retained, yielding a census of 28 eligible OMERO projects covering 284 datasets and 1765 image stacks [script “idr_scraping_full.py” and output file “idr_census.csv”, https://github.com/jbrocardplatim/FLATSTAT and *doi:10.5281/zenodo.20325392*]

FlatStat was subsequently applied to this census with additional eligibility filters: XY pixel size ≤ 10 µm, Z pixel size present and ≤ 10 µm, and maximum field dimension max(SizeX, SizeY) ≤ 4096 pixels. The final corpus comprised 1204 image stacks from 22 OMERO projects, yielding 4670 individual FlatStat measurements across all channels. All image data were retrieved on-the-fly via the OMERO API without local storage. [script “idr_flatstat_batch.py” and output file “flatstat_results.csv”, https://github.com/jbrocardplatim/FLATSTAT and *doi:10.5281/zenodo.20325392*]

### Bead acquisitions

To quantify the optical consequences of sample tilt, fluorescent bead slides were prepared and imaged on a Zeiss LSM800 confocal microscope (63× objective, NA 1.4). Tilt was induced at four conditions by adjusting the stage levelling screws; for each condition, planarity was measured using FlatStat (ImageJ macro implementation) on a stack of the Argolight SIM calibration slide (see Results, section 1). Then, three Z-stacks of multicolour fluorescent beads (1 µm multicolor TetraSpeck, ThermoFisher) were acquired per condition. Lateral and axial FWHM were measured on individual bead images with an ImageJ script and chromatic shift between green and red channels quantified using the MetroloJ_QC plugin [https://sites.imagej.net/MetroloJ_QC/plugins/] from ∼60 beads total per condition. All raw data and analysis scripts have been made available at https://github.com/jbrocardplatim/FLATSTAT and *doi:10.5281/zenodo.20325392*

### Data analysis

Slope distributions were summarized using the median as the central tendency measure, given their non-normal distribution. Between-project structuring of slope variability was quantified using η^2^ computed on log-transformed slopes (SS_between / SS_total), providing a descriptive measure of the proportion of total variance attributable to project-level differences [script “VAR_PROJECTS_LOG.R”, *doi:10.5281/zenodo.20325392*]

Inter-channel coherence was assessed in two complementary ways. Slope correlation between channel 0 and each additional channel within the same image was quantified by Spearman’s rank correlation. Directional coherence was assessed by cosine similarity between tilt vectors: cos = (vx_0_·vxc + vy_0_·vyc) / (|v_0_| × |vc|), where vx and vy are the Cartesian components of the tilt vector. Pairs where either vector had zero norm were excluded. [script “INTERCHANNEL_consistency.R”, *doi:10.5281/zenodo.20325392*]

## Results & Discussion

### 1. FlatStat: concept and first measurements

I developed FlatStat, an ImageJ macro that estimates sample tilt from any 3D fluorescence stack in three steps: a Z-map is computed by identifying, for each XY pixel, the focal plane of maximum intensity; a best-fit plane is then adjusted to the brightest pixels of this Z-map by least-squares regression; the resulting plane yields a mean slope in µm/100µm and a tilt direction expressed as a compass bearing (see Methods). To assess the planarity of the confocal system under investigation, I acquired images of two complementary samples: a geometrically defined Argolight SIM calibration slide, a spectrally stable reference free of biological variability and certified flat within 0.5 µm/100µm (Argolight, 2023) and a FluoCells prepared slide (Thermo Fisher Scientific) comprising two channels, a green-labeled actin network (C1) and a red-labeled mitochondrial network (C2), to test whether planarity could be estimated directly from a biological sample.

FlatStat was applied to four acquisitions of each sample (Figure 1). On the Argolight target, the four measurements yielded slopes of 0.51–0.56 µm/100µm (mean 0.55 ± 0.02 µm) with tilt directions of 332.2 ± 3.2°, a reproducibility that unambiguously attributed the measured tilt to the instrument rather than to acquisition noise or image content (Figure 1A). On FluoCells, actin network images showed comparable stability of mean slope (0.45 ± 0.03 µm, Figure 1B), while mitochondrial network images exhibited higher variability (0.55 ± 0.09 µm, Figure 1C), reflecting the influence of biological topography on the Z-map. Despite this increased dispersion in slope magnitude, tilt directions remained consistent across both channels and all four fields (347.7 ± 5.7°), confirming that the directional signal is instrument-driven rather than sample-specific.

**Figure 1.**
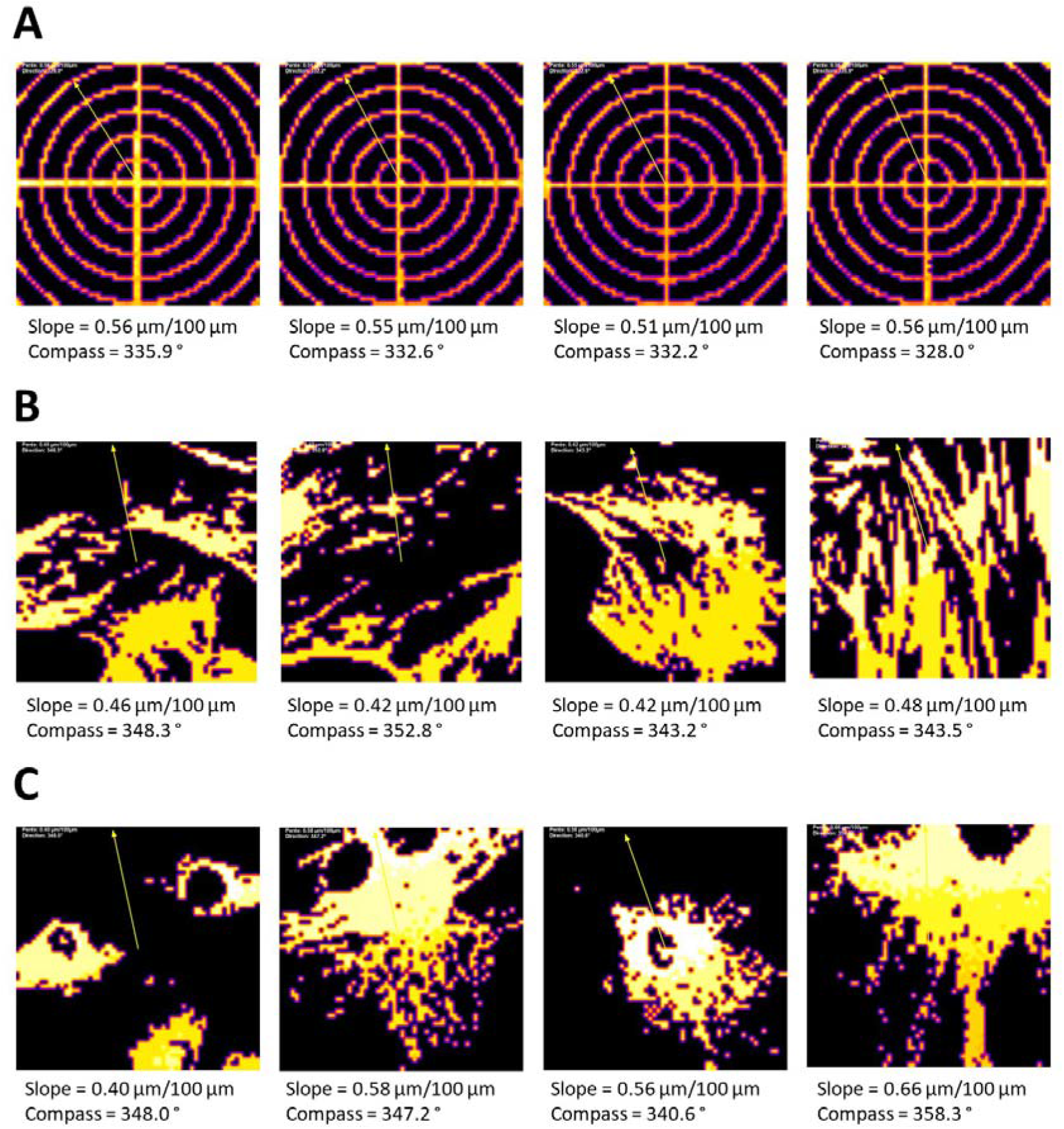
FlatStat measurements on two complementary sample types. **(A)** Four acquisitions of an Argolight SIM calibration slide (Argolight, Pessac, France). **(B)** Four fields of a FluoCells prepared slide (ThermoFisher), actin network. **(C)** Four fields as in B, mitochondrial network. For each image, the FlatStat Z-map is shown with the tilt vector superimposed (yellow arrow); slope in µm/100µm and compass direction are indicated below each panel.

These results established that FlatStat measurements on biological samples constitute a reliable proxy for the actual planarity experienced during imaging. The tilt direction measured on FluoCells (347.7 ± 5.7°) was offset by ∼ 15° relative to the Argolight target (332.2 ± 3.2°), across all four fields. Thus, it is probably attributable to the intrinsic geometry of coverslip mounting rather than to measurement noise. Indeed, FlatStat integrated contributions from stage tilt, slide geometry and sample flatness. The slope magnitude, meanwhile, was consistent across both sample types (0.50 ± 0.09 µm/100µm on FluoCells, 0.55 ± 0.02 µm/100µm on the Argolight target). Returning to the complaint that initiated this investigation: the user was right, a systematic focal plane deviation of 0.55 ± 0.02 µm/100µm was indeed present on this system. Whether this value was cause for concern, however, could not be answered without a reference distribution.

### 2. An opportunistic survey of community-scale planarity – corpus and slope distributions

No reference distribution of tilt measurements existed in the literature. However, the growing availability of FAIR fluorescence microscopy data in public repositories offered an unexpected opportunity to build one; IDR, built on the OMERO platform, proved uniquely suited to this purpose. Unlike generalist repositories, it allows direct programmatic retrieval of raw pixel data, making large-scale automated analysis practical. Of 144 IDR projects indexed at the time of the survey (March 2026), 28 met our initial technical eligibility criteria (see Methods). After additional pixel size, field dimension, and imaging method filtering, the final corpus comprised 1204 image stacks from these 22 OMERO projects, *i.e*. 4670 individual measurements across all channels (Table S1).

To enable large-scale analysis, FlatStat was reimplemented in Python, allowing on-the-fly slope and direction calculation from IDR image planes retrieved via the OMERO API (omeropy, Open Microscopy Environment, 2023) without local storage (see Methods). Applied to the final corpus, it yielded 4670 slope measurements across all fluorescence channels, each analyzed independently; channel 0, present in all 1204 stacks, served as the corpus-level reference, see representative examples Figure S1. Considering channel 0 only, slope distributions spanned several orders of magnitude between projects from 6–258 stacks per project, with a global mean of 1.78 ± 2.57 µm/100µm. The standard deviation exceeded the mean itself, reflecting extreme corpus heterogeneity (Figure 2A).

**Figure 2.**
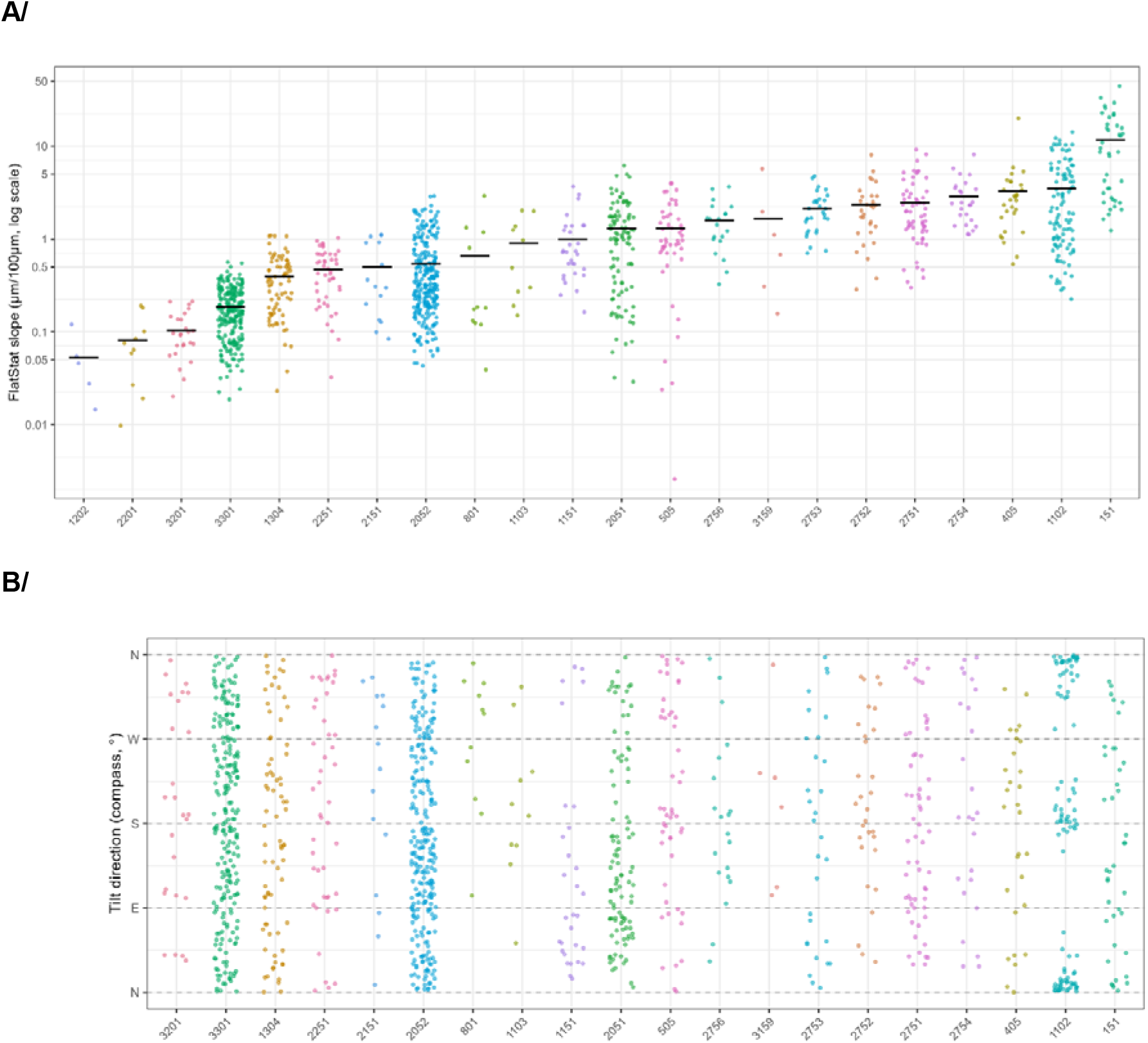
FlatStat slope and direction measurements across the IDR corpus (channel 0, n = 1204 image stacks). (A) Strip plot of slope measurements per project. Each point represents one image stack; horizontal bars indicate the median slope per project. The y-axis is on a log scale. Projects are ordered by increasing median slope; OMERO project IDs are used as labels on the x-axis. (B) Strip plot of raw tilt directions (dir_compass) by project, in the same order as in A. Each point represents one image stack. Dashed lines indicate the four cardinal directions (N = 0°/360°, E = 90°, S = 180°, W = 270°).

Indeed, close inspection revealed that two projects produced near-zero slopes and functioned as involuntary negative controls: #1202 (idr0085/MF-HREM block-face serial sectioning, median 0.046 µm/100µm), in which a mechanically flat surface was imaged by construction; and #2201 (idr0124/whole-embryo confocal imaging, median 0.070 µm/100µm), where whole embryos were imaged with no planar reference surface. In both cases, their consistently negligible slopes validated that FlatStat was not systematically measuring biological topography as tilt. As they did not represent conventional fluorescence microscopy of biological samples, they were excluded from all subsequent analyses. At the opposite extreme, project #151 (idr0027/chromatin imaging, median 8.66 µm/100µm) stood an order of magnitude above all other projects (Figure 2A). The associated publication (Dickerson et al., 2016) confirmed that these acquisitions involved thick 3D chromatin volumes imaged with large Z ranges, making high FlatStat slopes biologically plausible rather than artifactual. I therefore retained the project in the corpus as an informative outlier. As a more appropriate measure of central tendency, the median slope of the 20 remaining projects was measured at 1.28 µm/100µm, far exceeding the value measured on our own system (0.55 µm/100µm).

Slope variance was further decomposed: on log-transformed slopes, 58.8% of total variance was attributable to between-project differences, with 41.2% reflecting within-project variability (see Methods). As the aim was to characterize imaging systems rather than biological samples, image stacks from each OMERO project were treated as technical replicates of the same instrument configuration. A single publication may contribute several distinct projects to the corpus, each corresponding to a different experimental condition or acquisition mode — as illustrated by Hartmann et al. (eLife, 2020), whose projects #1102 (idr0079/experimentA, median 1.95 µm/100µm) and #1103 (idr0079/experimentB, median 0.72 µm/100µm) differed markedly despite originating from the same laboratory. This within-publication heterogeneity underscores that project-level treatment, rather than publication-level aggregation, is the appropriate unit of analysis for instrument characterization.

### 3. Tilt direction and inter-channel coherence

FlatStat computed a compass direction for every measured slope, regardless of its magnitude. Plotting these raw directions, across the 1187 channel-0 images with non-zero slopes from 20 projects, revealed a largely disordered landscape: most projects displayed directions scattered across the full 0–360° range, with no obvious preferred orientation (Figure 2B). One project, however, seemed to stand out: #1102 (idr0079/experimentA), where images clustered around the North–South axis. To quantify this signal, tilt directions were represented as rose diagrams for idr0079/experimentA alone and for the remaining corpus (Figure S2). The concentration around the NS axis in idr0079/experimentA was unambiguous (Rayleigh R = 0.41, p < 0.0001), while the remaining projects showed no detectable directional bias (Rayleigh R = 0.05, p = 0.096). This dataset represented lateral line tissue imaged in live conditions on a Zeiss LSM880 in AiryScan FAST mode (Hartmann et al., eLife 2020). Despite manual planarity correction documented in the original Methods section, FlatStat detected a strong and consistent NS orientation, more consistent with the acquisition mode than with biological or laboratory-dependent factors.

Furthermore, it was possible to use inter-channel coherence as an independent line of evidence on the origin of measured tilt. Across the 1187 channel-0 images, 1125 (94.8%) were acquired in more than one fluorescence channel, yielding 3404 channel-0/channel-n slope pairs. Slope measurements were strongly correlated between channels (ρ = 0.723, p < 2.2e-16), consistent with a shared physical origin — stage geometry or sample mounting — rather than channel-specific biological topography. Tilt direction consistency was also evaluated on these 3404 valid pairs: the median cosine similarity was 0.56 (Q1–Q3: −0.33 to 0.93), reflecting considerable dispersion. In project #1102 (idr0079/experimentA, see Figure S2), however, the strong NS bias was reflected with near-perfect inter-channel consistency (median cosine similarity = 0.94, Q1–Q3: 0.42–0.98, n = 43 pairs).

Surprisingly, project #151 (idr0027/chromatin imaging) combined the highest slopes in the corpus (median 8.66 µm/100µm, Figure 2A) with poorer inter-channel directional coherence (median cosine similarity = 0.66, Q1–Q3: −0.19 to 0.93, n = 40 pairs). This apparent dissociation suggests that the extreme slopes in idr0027 reflect genuine biological topography — thick chromatin volumes whose focal plane geometry varies between labellings — rather than a systematic instrumental bias (Dickerson et al., 2016). Taken together, these examples suggest that directional coherence, not slope magnitude alone, distinguishes instrumental from biological tilt.

### 4. Scarcity of instrument metadata limits causal inference

The strong directional signal identified in project #1102 (idr0079/experimentA) raised an obvious follow-up question: did AiryScan imaging systematically produce NS-oriented tilt across the IDR corpus, or was this observation confined to a single dataset? How would other imaging modes such as resonant scanner, spinning disk, two-photon, influence the directional signature of sample tilt and would systematic patterns emerge across instruments and laboratories? Answering these questions would require identifying the acquisition mode for each image in the corpus, which in turn requires reliable instrument metadata.

Structured OMERO instrument objects (microscope model, objective, detector) were present for approximately 16% of images in the census, covering 5 of the 22 eligible OMERO projects. Free-text annotations contained instrument-related information for only a small minority of projects, in a heterogeneous and unparseable form: microscope models appeared embedded in protocol descriptions, as manufacturer trade names, or not at all. No dataset-level instrument metadata was available in any of the 22 projects analyzed. In practice, the acquisition mode responsible for the NS bias in idr0079/experimentA could only be identified because the original publication (Hartmann et al., eLife 2020) explicitly described it, not from the IDR metadata.

This reflected a community-wide absence of standardized instrument metadata at the point of data deposition. Initiatives such as REMBI and the Quarep-LiMi working group have begun to address this gap, but adoption remains limited (Sarkans et al., 2021; Nelson et al., 2021). From the perspective of FlatStat, the practical consequence was clear: planarity measurements can be made at scale, but their interpretation cannot be systematically resolved without knowing what instrument produced the data. Beyond instrument metadata, the within-project slope variability (41.2% of total variance) suggested that sample preparation factors - including slide lot, mounting medium, and operator – also contributed substantially to tilt, independently of the imaging system. Hence, both instrument and sample preparation metadata could be considered as structured, mandatory fields at the point of data deposition, a step that would transform an observational survey of this kind into a genuinely explanatory study (see Bajcsy et al., 2025).

### 5. Optical consequences of planarity defects: back to the original complaint

The IDR survey established that sample tilt was both common and instrumentally structured across the fluorescence microscopy community. Whether it actually mattered for image quality, however, remained an open question, and one that brought me back to the confocal system whose focal plane had prompted a user complaint.

To quantify the optical consequences of tilt, I induced four tilt conditions on the stage of our system by adjusting the levelling screws. Each condition was verified on an Argolight SIM calibration slide using FlatStat (0.28 to 3.14 µm/100µm, Figure 3A) and three fields of multicolor 1-µm fluorescent beads were acquired on two channels per condition. From these, 3D chromatic shift was measured using the MetroloJ_QC plugin of FIJI, while lateral and axial resolutions were approximated by FWHM measurements with a homemade FIJI script (see Methods).

**Figure 3.**
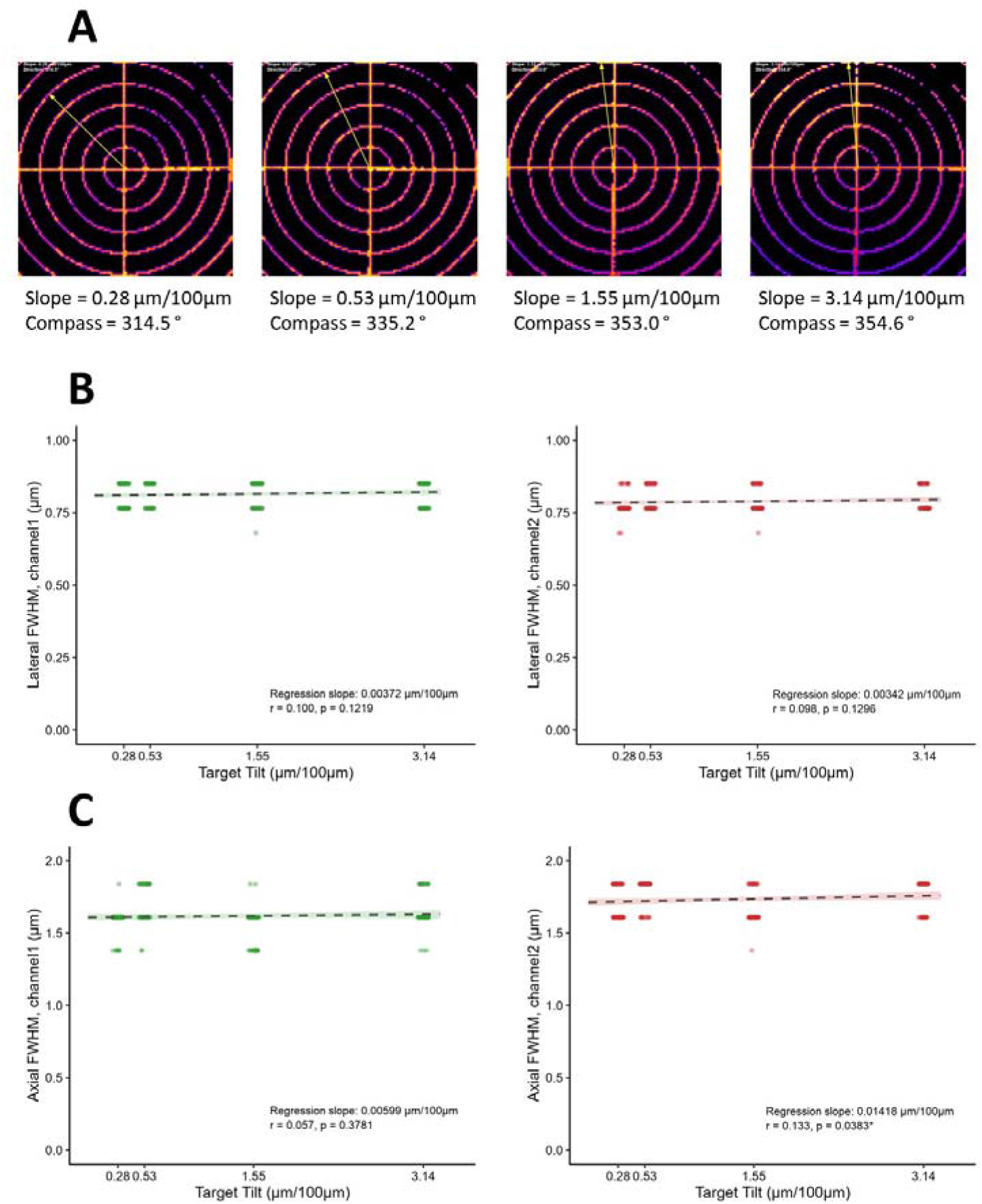
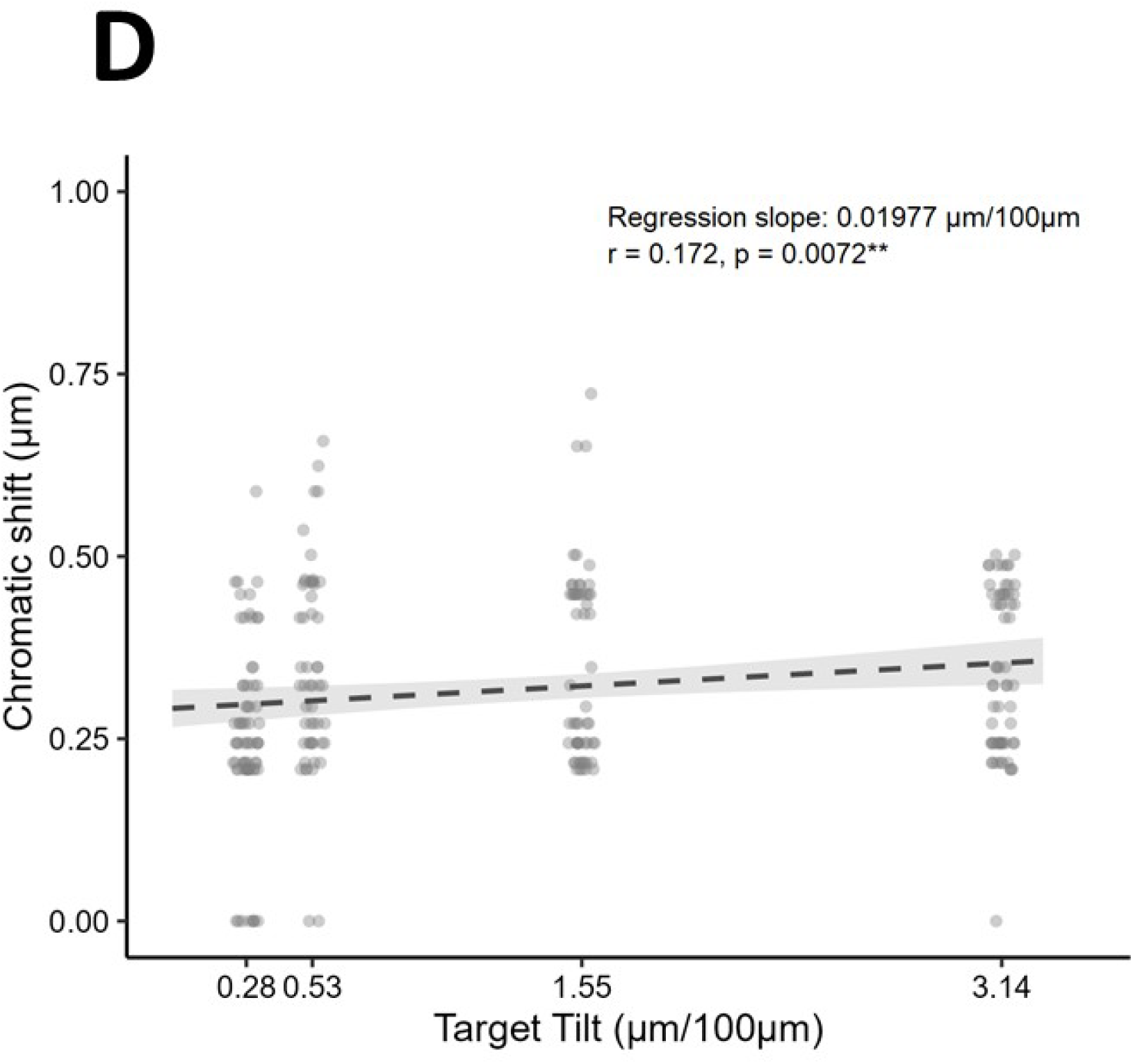
Optical consequences of sample tilt on lateral resolution, axial resolution and chromatic shift. (A) FlatStat Z-maps of the Argolight target at each tilt condition, with the measured slope (µm/100µm) and compass direction indicated below each panel. (B) Lateral FWHM in channel 1 (green, left) and channel 2 (red, right) measured on fluorescent beads (n = 69, 53, 57 and 63 beads at 0.28, 0.53, 1.55 and 3.14 µm/100µm respectively). (C) Axial FWHM in channel 1 (green, left) and channel 2 (red, right) measured on the same fluorescent beads as in B. (D) 3D chromatic shift measured between channels 1 and 2. In panels B–D, individual bead measurements are shown as dots and the dashed line represents the linear regression fit with its 95% confidence interval (shaded area), regardless of significance. Regression slope, Pearson r and p-value are indicated in each panel. *, p<0.05; **, p<0.01

Lateral resolution was entirely insensitive to tilt across the full range tested (channel 1: r = 0.100, p = 0.1219; channel 2: r = 0.098, p = 0.1296; Figure 3B). Axial resolution showed a weak but nominally significant trend in channel 2 only (r = 0.133, p = 0.0383), with no detectable effect in channel 1 (r = 0.057, p = 0.3781, Figure 3C). Finally, chromatic shift showed the clearest response, increasing significantly with tilt (r = 0.172, p = 0.0072; Figure 3D), with a regression slope of 0.0198 µm per µm/100µm of tilt, corresponding to a net increase of approximately 57 nm over the full tilt range tested.

So, was the user right to worry? In principle, yes: a systematic tilt of 0.55 µm/100µm is real, measurable, and did produce a slight modification of measured parameters. In practice, however, the effect would be modest: chromatic shift amounted to ∼0.30 µm at the lowest induced tilt (0.28 µm/100µm), increasing to ∼0.31 µm at the time of the complaint, a difference unlikely to affect any experiment in photonic microscopy.

## Conclusion

This study set out to answer a deceptively simple question: how flat are fluorescence microscopy samples, and does it matter? FlatStat provided a practical, automated answer to the first part, deployable on any 3D fluorescence stack, without prior knowledge of sample content, in two complementary implementations (ImageJ macro and Python script). Applied opportunistically to 1204 image stacks from 22 IDR projects, it revealed that sample tilt is ubiquitous, strongly structured at the instrument or laboratory level, and carried a clear directional signature in one specific instance, reflecting the acquisition mode rather than the biology.

The answer to the second part is more nuanced. On the single system examined here, tilt within the range commonly observed in the community produced measurable but very modest optical consequences. Whether these consequences cross a biologically relevant threshold will depend on the system, the objective correction, and the application.

Establishing community reference values across instruments and modalities is precisely the type of multi-site, standardized campaign that the Quarep-LiMi consortium is positioned to undertake, with FlatStat providing the planarity measurement component.

## Supporting information

FLATSTAT_v6_supplementary

## Acknowledgments

I acknowledge the unwitting contribution of A. Basile (Laboratoire de Biologie et Modélisation de la Cellule, UMR5239, ENS de Lyon, France), whose observations prompted this investigation. This study was carried out at the imaging core facility LyMIC-PLATIM of the SFR Biosciences (Université Claude Bernard Lyon 1, CNRS UAR3444, Inserm US8, ENS de Lyon, France) and I thank my colleagues there for stimulating discussions throughout: Ema Bobocioiu-Caracas, Elodie Chatre and Jean-Luc Duteyrat. This work was supported by the National Infrastructure France-BioImaging funded by the French National Research Agency (ANR-24-INBS-0005 FBI BIOGEN).

## References

Argolight, 2023: https://argolight.com/files/Argo-SIM/Argo-SIM-v1.1_User-guide.pdf

Bajcsy P, Bhattiprolu S, Börner K, Cimini BA, Collinson L, Ellenberg J, et al. Enabling global image data sharing in the life sciences. Nature Methods, 2025, 22(4):672–676. doi: 10.1038/s41592-024-02585-z

Dickerson D, Gierliński M, Singh V, Kitamura E, Ball G, Bhalla N, Owen-Hughes T. High resolution imaging reveals heterogeneity in chromatin states between cells that is not inherited through cell division. BMC Cell Biology,2016, 17:33. doi: 10.1186/s12860-016-0111-y

Faklaris O, Bancel-Vallée L, Dauphin A, et al. Quality assessment in light microscopy for routine use through simple tools and robust metrics. Journal of Cell Biology, 2022, 221(5):e202107093. doi: 10.1083/jcb.202107093

Hartmann J, Wong M, Gallo E, Gilmour D. An image-based data-driven analysis of cellular architecture in a developing tissue. eLife, 2020, 9:e55913. doi: 10.7554/eLife.55913

Le Bourdellès G, Mercier L, Roos J, Bancelin S, Nägerl UV. Impact of a tilted coverslip on two-photon and STED microscopy. Biomedical Optics Express, 2024, 15(2):743–752. doi: 10.1364/BOE.510512

Nelson G, Boehm U, Bagley S, Bajcsy P, Bischof J, Brown CM, Dauphin A, Dobbie IM, Eriksson JE, Faklaris O, et al. QUAREP-LiMi: A community-driven initiative to establish guidelines for quality assessment and reproducibility for instruments and images in light microscopy. Journal of Microscopy, 2021, 284(1):56–73. doi: 10.1111/jmi.13041

Open Microscopy Environment, 2023: https://pypi.org/project/omero-py/

Sarkans U, Chiu W, Collinson L, et al. REMBI: Recommended Metadata for Biological Images — enabling reuse of microscopy data in biology. Nature Methods, 2021, 18:1418–1422. doi: 10.1038/s41592-021-01166-8

Williams E, Moore J, Li S et al. Image Data Resource: a bioimage data integration and publication platform. Nature Methods, 2017, 14(8):775–781. doi: 10.1038/nmeth.4326

