## Supplementary material for "How flat is your sample? An opportunistic survey of 3D tilt in public fluorescence microscopy data": FLATSTAT_v6_supplementary

**SUPPLEMENTARY FIGURES**

**Figure S1. Representative FlatStat Z-maps from the IDR corpus.** Each panel shows the Z-map computed by FlatStat for one image stack from the indicated OMERO project (project ID shown in the top-left corner of each panel). The colour scale indicates the Z position of maximum intensity (µm); the white arrow indicates the tilt direction and is proportional to the slope magnitude. Measured slope (µm/100µm) and compass direction are indicated in each panel title. The five white crosses indicate the tilt vector estimates from percentile thresholds 88–92, prior to averaging (see Methods).

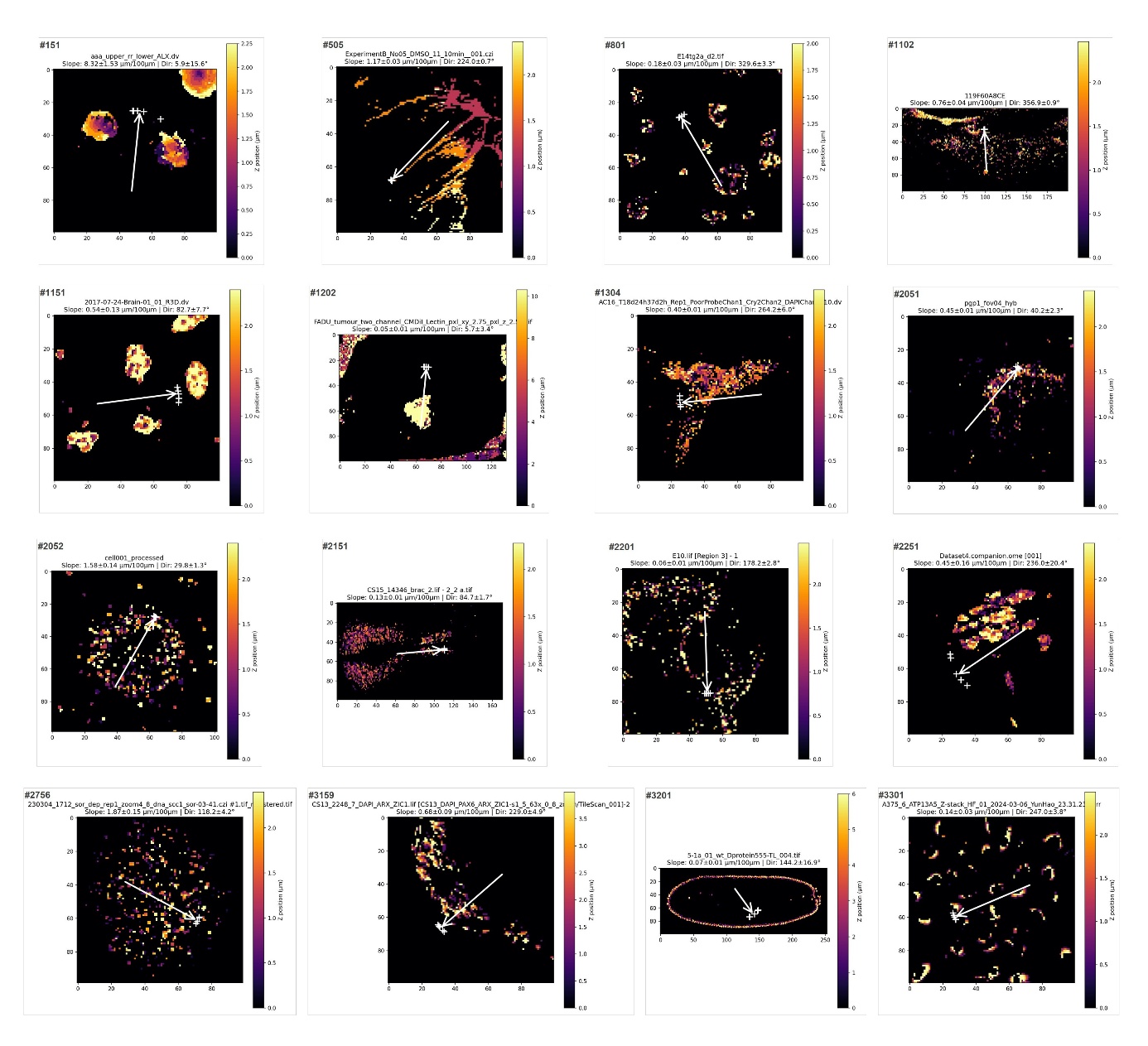

**Figure S2.** **Rose diagrams of tilt directions by project subset (channel 0).** (A) Tilt directions for project #1102 (idr0079/experimentA, n = 100 image stacks). The distribution is strongly concentrated along the North–South axis (max section = 33 hits); the red arrow indicates the mean resultant vector (Rayleigh R = 0.41, p < 0.0001). (B) Tilt directions for all remaining projects (n = 1087 image stacks). The distribution shows no detectable directional bias (max section = 86 hits; Rayleigh R = 0.05, p = 0.096).

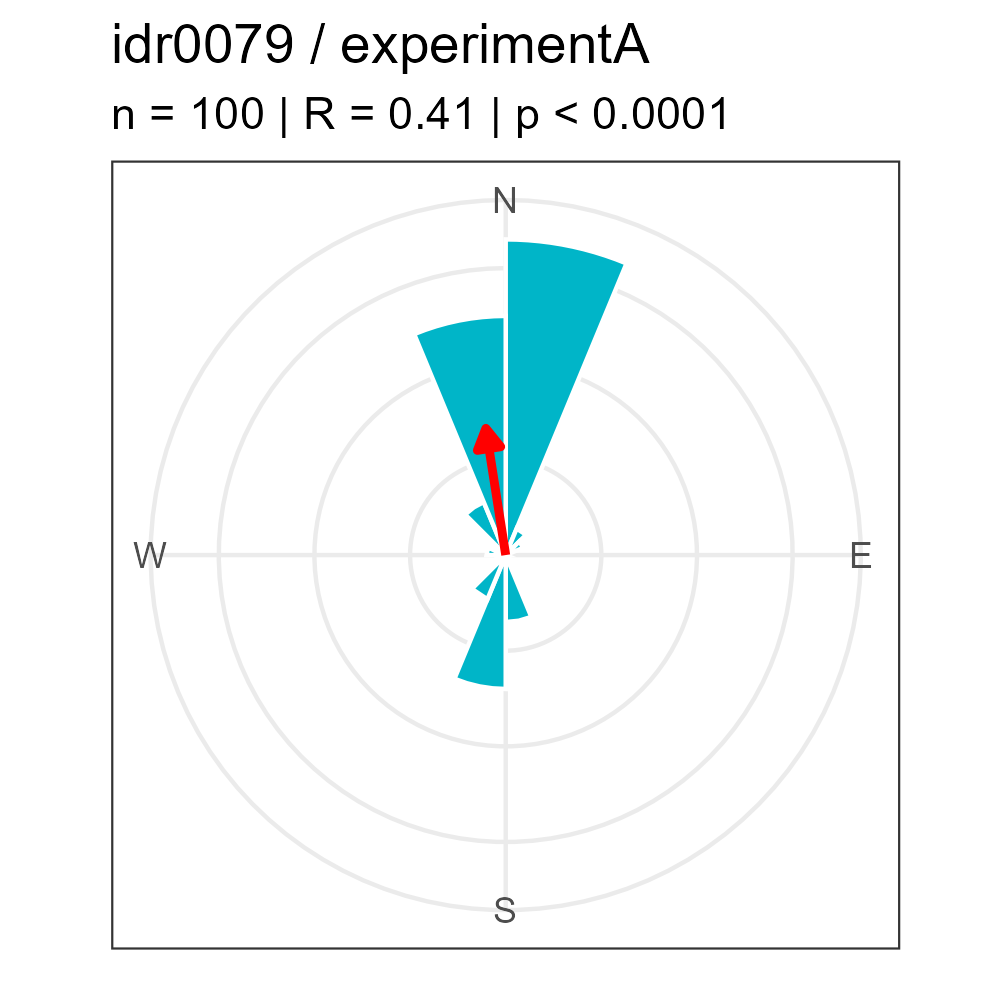

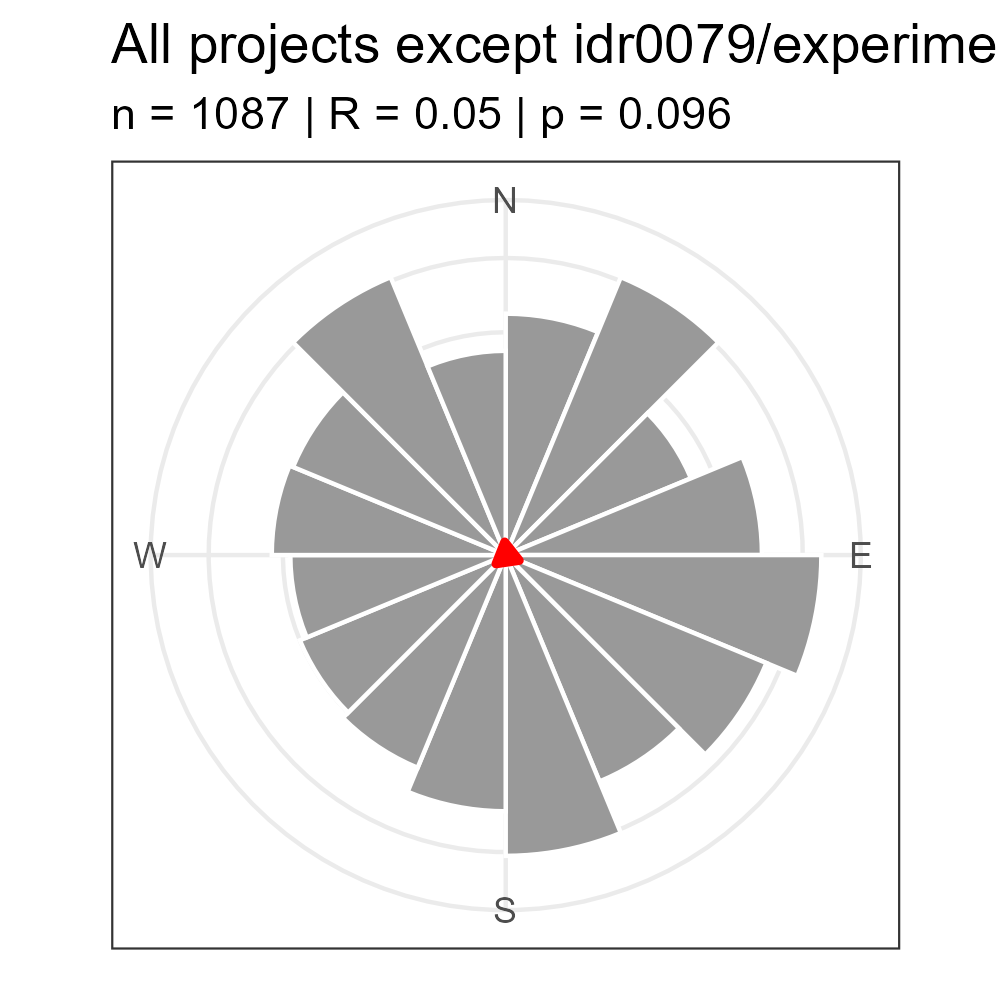

**Table S1. Summary of FlatStat measurements across the IDR corpus.** For each OMERO project retained in the analysis, the table reports the project name and identifier (project name, project id), the number of image stacks (#stacks) and mean channel measurements (#channels), and the median and mean slope of tilt in µm/100µm measured in channel 0 (med_slope_c0 and mean_slope_c0). Projects are ordered by decreasing median slope. *The two negative control projects (idr0085, idr0124) are shown separately at the bottom of the table.*

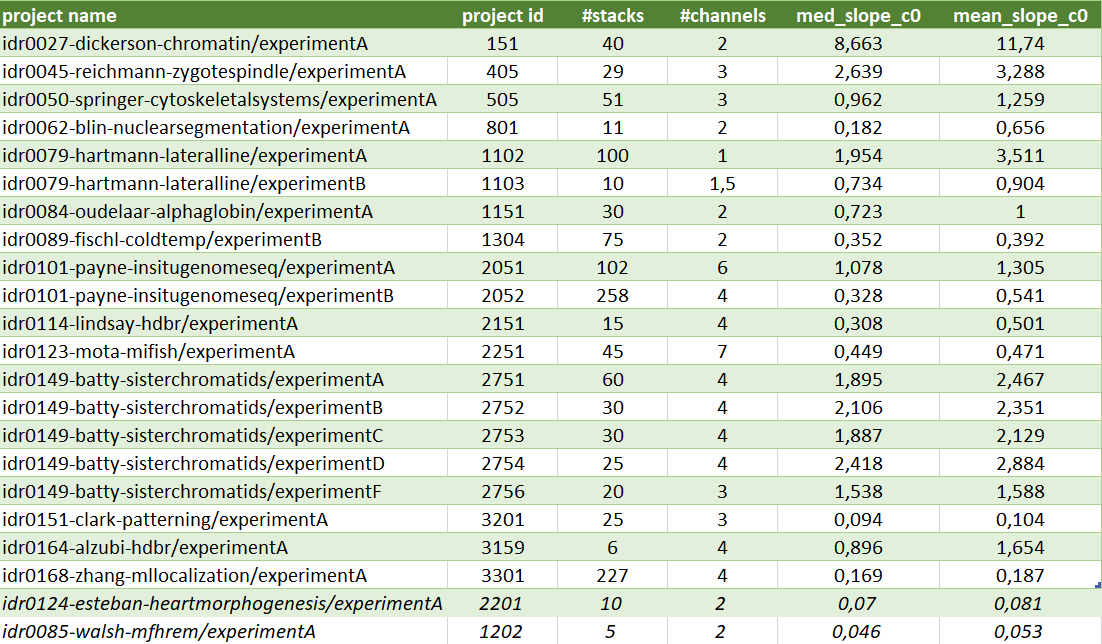
